# SNAPPY: <u>S</u>ingle <u>N</u>ucleotide <u>A</u>ssignment of <u>P</u>hylogenetic <u>P</u>arameters on the <u>Y</u> chromosome

**DOI:** 10.1101/454736

**Authors:** Alissa L. Severson, Jonathan A. Shortt, Fernando L. Mendez, Genevieve L. Wojcik, Carlos D. Bustamante, Christopher R. Gignoux

## Abstract

**Summary:** The assignment of Y chromosome data to related clusters, or haplogroups, is a common application in human population genetics. To enable this at scale, we developed SNAPPY. SNAPPY is a software program used to assign Y-chromosome phylogeny-informed haplotypes using dense genotype data. The program efficiently tests all haplotypes in a provided Y-chromosome database to find the haplogroup that is best supported by the input genotypes. Importantly, the method considers both the amount of support for the specific haplogroup, as well as its ancestral haplogroups via parsimony. This accounts for the underlying genealogy the haplotypes represent, strengthening the accuracy of the assignments. SNAPPY is fast, scalable, and uses standard file formats, making it easy to integrate into analytical pipelines.

**Availability and Implementation:** The program is implemented in python. The program, a user manual, haplotype databases, and test datasets are available for download at github.com/chrisgene/snappy.

## Introduction

Analyses of Y-chromosome haplogroups have yielded important insights into human migration history (Bergstrom, et al., 2016; Chiaroni, et al., 2009), cultural customs (Seielstad, et al., 1998), and ancestral population sizes (Karmin, et al., 2015). These insights are possible because of the two unique characteristics of the Y-chromosome: it is passed directly from father to son, and much of the Y-chromosome does not recombine with any other chromosome; this leaves a long tract of unbroken DNA that serves as faithful record of evolution of the chromosome throughout human history. Constant accumulation of variation within the chromosome over human history created new combinations of variants (haplotypes), which trace patrilineal descent.

Despite its use in ancestry, efficient software tools that are able to rapidly assign Y-chromosome haplotypes from dense genotype data into clusters of related haplotypes (haplogroups) at population scales are limited in scope. Here we describe SNAPPY (<u>S</u>ingle <u>N</u>ucleotide <u>A</u>ssignment of <u>P</u>hylogenetic <u>P</u>arameters on the <u>Y</u>-chromosome), a tool to assign Y-chromosome haplogroups using dense genotyping data. Haplogroup assignment is based on the well-established polymorphisms along the Y-chromosome with haplogroup information maintained by the International Society of Genetic Genealogists (ISOGG, isogg.org) and others [e.g., (Karafet, et al., 2015; Poznik, et al., 2016)], using only phylogenetically-informative alleles to determine which haplogroup has the highest support, thus avoiding complications related to the reversion of alleles to ancestral states. Importantly, the method is able to identify haplogroups from both leaf and interior nodes of the Y-chromosome tree. Here we briefly outline the implementation of the program, and present Y-chromosome haplotype assignments from The 1000 Genomes Project (Genomes Project, et al., 2015) data to validate the algorithm.

## Implementation

### Dependencies

SNAPPY is implemented in python requiring only numpy in addition to the standard python library. In addition, SNAPPY makes use of plink (Chang, et al., 2015) for initial file format conversions from either array-based formats (e.g., .ped/.bed) or from vcf.

### Algorithm Description

SNAPPY leverages the Y-chromosome phylogeny and a database of haplotype-informative SNPs derived from ISOGG and other sources to store nested haplotypes in memory-efficient dictionaries. Genotypes from individual samples are stored as a list of dictionaries keyed by chromosome position. Currently the tree is optimized for sites on the commonly-used Illumina Multi-Ethnic Global Array (pagestudy.org/mega), however tree files can easily be generated from relevant sources. To compute haplogroup scores for each of the 565 possible Y-chromosome haplogroups present in our Y-chromosome haplogroup reference library, SNAPPY looks up a sample’s genotypes—stored in a dictionary—and counts the proportion of matching alleles that are at haplogroup-informative positions.

SNAPPY determines a score for each haplotype using the number of haplogroup-informative derived alleles adjusted by the sample’s number of non-missing informative positions. Note that a particular haplogroup’s score uses alleles from both its own informative positions as well as its ancestral nodes on the tree. This ensures that haplogroup assignments take into account the full phylogenetic structure of the Y-chromosome, and enables the creation of easily traversable data structures that eliminate redundant storage of informative positions due to the highly parsimonious nature of variants on the Y(Poznik, et al., 2016). Haplogroup scores for every haplogroup are stored in a two-dimensional numpy array to allow for efficient storage and quick processing.

Finally, haplogroup assignments are made by evaluating the scores of nodes of each branch, starting at the most distal node of the branch that has both a score above a user-defined threshold and no descendant haplotypes with a score that exceeds the user-defined threshold, and then working towards the root of the tree. This ensures that all nodes are considered potential haplogroups for the sample, even if they are not terminal nodes on the tree. SNAPPY makes its haplogroup assignment based on the highest scoring node or the deepest node with a score higher than a user-defined threshold. This algorithm ensures that the deepest haplogroup with sufficient support is assigned. In addition to reporting the single haplogroup with the most support for each sample, SNAPPY also reports each haplogroup that has a score greater than a user-defined minimum score so that haplogroup assignments can be adjusted or investigated where necessary.

## Validation and Testing

### Data Sources

We downloaded a list of Y-chromosome variants found on the Multi-Ethnic Genotyping Array (MEGA) (Bien, et al., 2016) (a list of variants found on the MEGA can be found at https://pagestudy.org/index.php/multi-ethnic-genotyping-array). Because the MEGA Y content was designed to be ancestry informative, the SNPs included are well-distributed throughout the Y-chromosome tree, which allows for accurate and precise assignment of haplogroups. MEGA variant positions were converted from GRCh38.p7 to GRCh37 using NCBI’s genome remapper tool (https://www.ncbi.nlm.nih.gov/genome/tools/remap) in preparation for extracting MEGA positions from Y-chromosome Phase 3 of The 1000 Genomes Project (Genomes Project, et al., 2015) data.

For reference, we downloaded 1,233 Y-chromosome genotypes from Phase 3 of The 1000 Genomes Project from NCBI (Genomes Project, et al., 2015) (ftp://ftp-trace.ncbi.nih.gov/1000genomes/ftp). The vcf was filtered to contain only the variants that are present on MEGA using plink (Chang, et al., 2015). This yielded a final total of 2,366 variants.

Similarly, we provide Y-chromosome haplogroup tree information and informative SNPs, available in the SNAPPY distribution found at github.com/chrisgene/snappy. This information is largely consistent with phylogenetic information present on ISOGG.

### Results

To assess the speed and accuracy of SNAPPY, we tested 1,233 males from Phase 3 of The 1000 Genomes Project (Genomes Project, et al., 2015). The program is computationally efficient: for a set of 1,233 samples genotyped at 2,366 SNPs, it runs in an average of 4.96 seconds on a single 2.3 GHz core of an Intel Xeon processor, using 278 Mb of RAM. SNAPPY scales linearly indicating good performance for larger data sets.

Y-chromosome haplotypes for 1000 Genomes males have been previously well-characterized (Poznik, et al., 2016), and we used this characterization to assess the accuracy of SNAPPY on genotype data compared to full sequences. We found that most individuals had top haplogroup scores >95%, (Supplemental Figure 1), correctly predicting over >99% of major haplogroup assignments for all individuals, with minor differences in fine-grained haplogroup designations, even given topological differences between our tree and prior examples (Supplemental Table 1). The three individuals with discordant major haplogroup assignments were assigned by SNAPPY to the haplogroup P1 rather than the closely-related reference assignment of Q1a. We note that the P1 and Q haplogroups had the same score (0.958) in these individuals, but the P1 haplogroup was chosen because it resides deeper in the tree than Q. These inconsistencies are expected to resolve with increased genotype density or tree topologies with improved resolution and can be easily adjudicated on manual inspection. A fourth individual was correctly assigned to the A0 haplogroup consistent with no derived haplogroup with support above our recommended 60% minimum match. Some differences between the sets are to be expected because the number of variants between the two datasets differ substantially (>60K for the reference set vs. 2,399 for our tests), and different Y-chromosome phylogenies (*de novo* reconstruction vs. ISOGG download) may result in subtle yet important differences between branch points. Nevertheless, the ability to recover major haplogroups from individuals representing nearly all major branches of the Y haplogroup tree indicates that SNAPPY, and informative array-genotyped sites, are robust and unbiased in ancestry assignment.

## Conclusions

The ability to determine Y-chromosome haplogroups is of broad interest in many lines of research from population genetics to anthropology to medicine. Here we have introduced and demonstrated SNAPPY, a method to rapidly and accurately assign Y-chromosome haplogroups to population-scale data sets using dense genotyping. SNAPPY is user-friendly, requiring input in common plink format and just a single command to run. Additionally, SNAPPY is flexible, with several user-defined parameters, the ability to use user-curated y-haplogroup trees, and to manually interpret haplogroup assignments.

## Acknowledgments

We gratefully acknowledge the community in ISOGG for maintaining and providing Y-chromosome haplogroup trees, David Poznik for assistance with the original phylogeny, and Peter Underhill for general discussions.

---

**Supplemental Figure 1.**
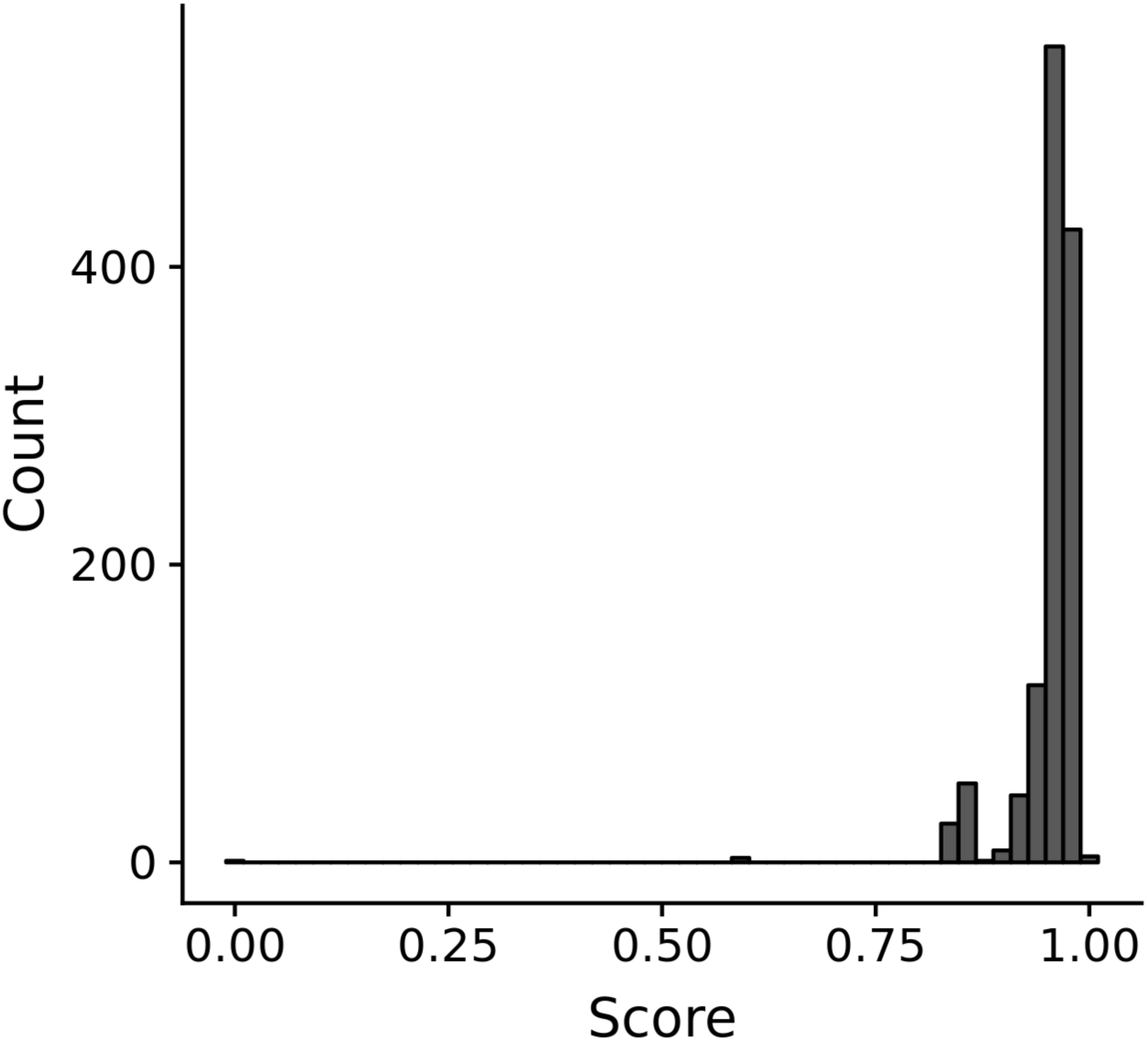
Distribution of SNAPPY-assigned haplogroup scores. Histogram of haplogroup scores assigned by SNAPPY. A dashed red line indicates 95%.

**Supplemental Table 1.**
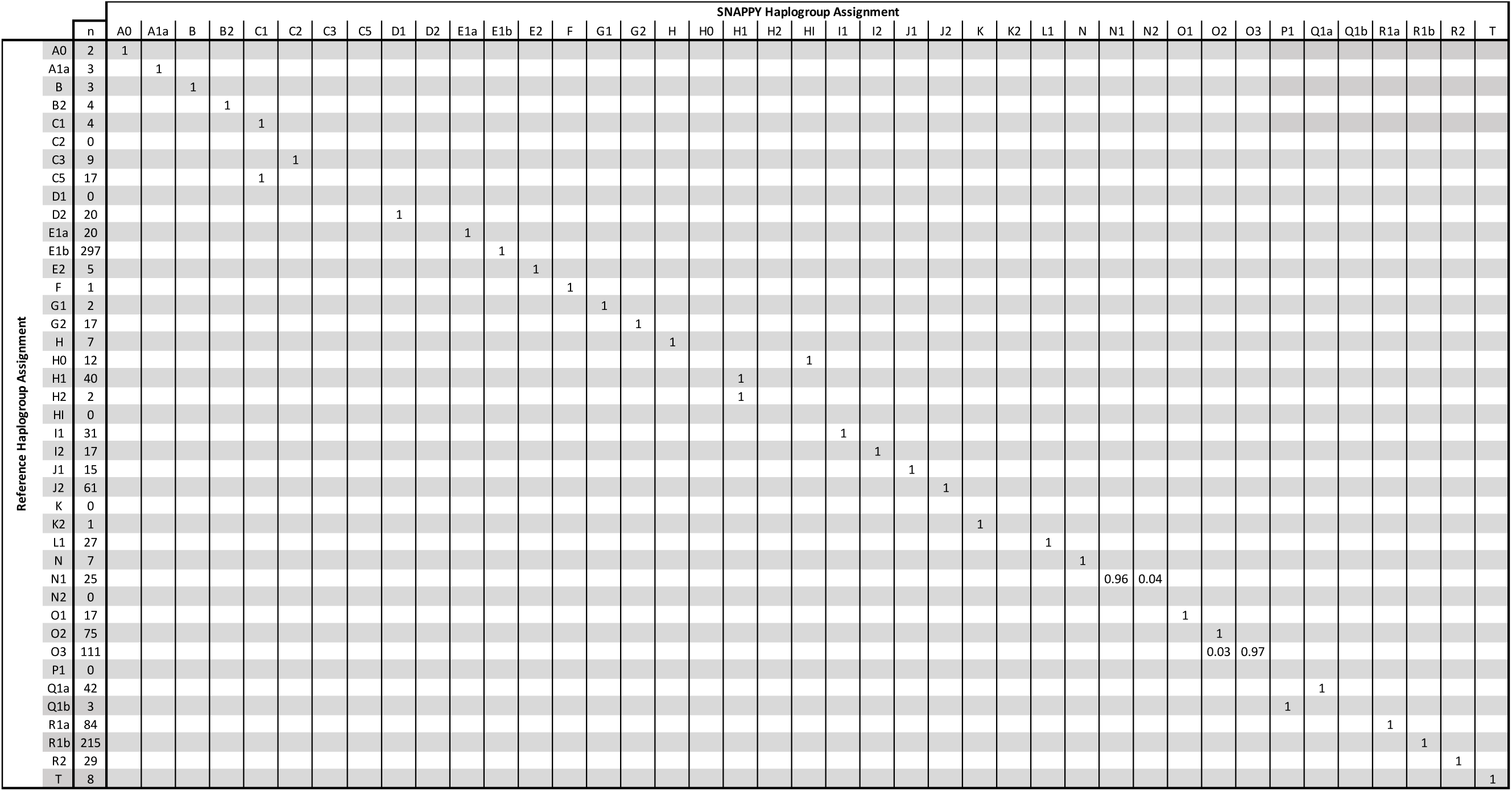
Proportion of Concordant and Discordant SNAPPY Haplotype Assignments Relative to Major Reference Haplotypes Johnson et al. 1

